# Home range and dynamic space use reveals age-related differences in risk exposure for reintroduced parrots

**DOI:** 10.1101/2022.12.17.520898

**Authors:** S. W. Forrest, M. Rodríguez-Recio, P. J. Seddon

## Abstract

Individual-level differences in animal spatial behaviour can lead to differential exposure to risk. We assessed the risk-exposure of a reintroduced population of kākā (*Nestor meridionalis*) in a fenced reserve in New Zealand by GPS tracking 10 individuals and comparing the proportion of each individual’s home range beyond the reserve’s fence in relation to age, sex, and fledging origin. To estimate dynamic space use, we used a sweeping window framework to estimate occurrence distributions from temporally overlapping snapshots. For each occurrence distribution, we calculated the proportion outside the reserve’s fence to assess temporal risk exposure, and the area, centroid and overlap to represent the behavioural pattern of space use. Home range area declined significantly and consistently with age, and the space use of juvenile kākā was more dynamic, particularly in relation to positional changes of space use. The wider-ranging and more dynamic behaviour of younger kākā resulted in consistently more time spent outside the reserve, which aligned with a higher number of incidental mortality observations. Quantifying both home range and dynamic space use is an effective approach to assess risk exposure, which can provide guidance for management interventions. We also emphasise the dynamic space use approach, which is flexible and can provide numerous insights towards a species’ spatial ecology.

## Introduction

Individual and group-level behavioural differences have been observed in a broad range of animal taxa, which represents intrapopulation diversity in physiological, behavioural, and ecological mechanisms (Bolnick et al., 2003; Dall et al., 2012; Stuber et al., 2022). These differences can manifest as variable foraging strategies, spatial behaviours (Stuber et al., 2022), and problem-solving abilities of conspecific individuals (Loepelt et al., 2016), each of which influence population dynamics (Hanski, 1998). Extrinsic factors that influence behaviour include resource availability and the density of competitors, predators, prey or conspecifics, while intrinsic factors include age, sex, personality and breeding status (Andreassen et al., 2002; Class et al., 2019; Clay et al., 2018; Lesmerises & St-Laurent, 2017; Loepelt et al., 2016; Reader & Laland, 2002; Stuber et al., 2022). In birds, age is considered to influence behaviours such as those relating to problem solving and foraging efficiency, with younger individuals being more behaviourally flexible, innovative, and able to solve novel problems (Loepelt et al., 2016; Sherratt & Morand-Ferron, 2018) whilst older individuals are typically more efficient foragers (Clay et al., 2018; Phillips et al., 2017). Age-related plasticity is evidenced by changes in behaviour in relation to age, when there is not selective disappearance (Class et al., 2019).

In a conservation context, individual- or group-level behavioural differences that lead to variability in space use may lead to differential exposure to risk, which can affect the population’s or species’ trajectory and persistence (Merrick & Koprowski, 2017). In the case of fully-fenced conservation reserves, which we consider to be discrete areas of habitat that undergo protection from threats and allow for the refuge and possible translocation of sensitive wildlife species that have been reduced or historically extirpated, even while the major causes of species decline persist in the wider region (Burns et al., 2012; Innes et al., 2019), flighted species that ‘spill-over’ into the wider landscape (Tanentzap & Lloyd, 2017) may encounter a fragmented or modified landscape containing a higher number of threats. The exposure to threats can impose a greater mortality risk outside the fence, which may be further increased if the species has lost predator-avoidance behaviours (Muralidhar et al., 2019).

Differences in spatial behaviours that lead to variable exposure to risk can be assessed through telemetry data and space use analyses. Range distributions are a probabilistic representation of the space required by an animal over the long term (i.e. its home range), and occurrence distributions are a probabilistic representation of the space used by an animal during a defined period of observation (Alston et al., 2022; Fleming et al., 2015, 2016). We can therefore estimate long-term risk exposure by calculating the overlap of an individual’s home range with areas of presumed risk, or risk exposure during a specified time period using an occurrence distribution (Alston et al., 2022). While the home range approach provides valuable evidence and can indicate whether certain individuals or groups are likely to be at higher risk than others over long time scales, an animal’s space use may not be stable, and resources or climate might drive the space use of certain individuals, groups, or even the entire population to change in position or size, leading to risk-exposure that changes dynamically. Estimating space use dynamically using overlapping occurrence distributions can therefore provide information about the pattern of space use, and whether risk-exposure is likely to change throughout time. The term utilisation distribution (UD) is often used to describe both range and occurrence distributions (Alston et al., 2022; Fleming et al., 2015), although to minimise ambiguity here we differentiate by using the term home range (HR) to denote a range distribution and OD to denote an occurrence distribution.

We aimed to assess risk exposure using range distributions and dynamic space use in a reintroduced parrot population in a fenced conservation reserve, and to highlight the utility of both approaches. Kākā (*Nestor meridionalis*) are large-brained, social parrots (Bond & Diamond, 2004; Iwaniuk et al., 2005) that are considered generalist, extractive foragers (Beggs & Wilson, 1987; Moorhouse, 1997). These traits suggest a propensity for exploration and innovation (Dunbar & Shultz, 2007; Sol et al., 2005), which has been observed particularly in juvenile kākā (Bond & Diamond, 2004; Loepelt et al., 2016). Threats to kākā in wild populations are well-understood, which are predominately the destruction of native habitat, predation by stoats (*Mustela erminea*) and competition for food resources by vespulid wasps and possums (*Trichosurus vulpecula*) (Beggs & Wilson, 1991; Greene et al., 2004; R. Moorhouse et al., 2003; O’Donnell & Rasch, 1991; Wilson et al., 1998). However, fragmented, semi-urban landscapes present novel threats that include human infrastructure, vehicles, cats, dogs, and traps and toxins intended for pest species. In 2008, kākā were translocated into the fully fenced semi-urban conservation reserve Orokonui Ecosanctuary (*Te Korowai o Mihiwaka*; 45°46 ’S, 170°36 ’E; hereafter Orokonui) in New Zealand, which is within a region that kākā were absent from for many years. We recognised the continued presence of threats outside the reserve and hence attempted to understand to what extent movements outside the apparent safety of the fenced reserve pose a risk to the birds, and whether this differs between certain groups. To assess whether analyses of home range and dynamic space use can address these questions, we GPS-tracked a subset of the kākā of the Orokonui population. We assessed the kākā’s exposure to risk by estimating the proportion of their home range outside the Orokonui Ecosanctuary fence and described behavioural patterns by estimating their dynamic space use. The approach taken here has broad application for quantifying exposure to presumed risk and when this occurs, such as outside conservation reserves, within fishing grounds for marine species (Weinstein et al., 2017), and in areas with human infrastructure such as wind turbines (Bright et al., 2008).

## Methods

### Ethical note

This study was conducted with approval from the University of Otago Animal Ethics Committee under the Animal Use Protocol AUP-18-237. We consulted with the Ngāi Tahu Research Consultation Committee and Kati Huirapa Runaka ki Puketeraki prior to undertaking the research, and all kākā handling was undertaken by personnel trained by Department of Conservation staff with Level 3 bird banding permits. Free-ranging kākā were captured in the reserve for tagging by luring them into a baited aviary used for soft-release of captive-raised individuals during population reinforcement translocations. All kākā were weighed, subjected to a health check, and sexed by measuring the upper mandible (Moorhouse & Greene, 1995; Moorhouse et al., 2008). GPS devices were attached to kākā using backpack harnesses with a weak-link (Karl & Clout, 1987), and we followed standard procedures for attaching devices and reduced handling time as much as possible to minimise stress to the kākā. Nine of the GPS devices (Lotek PinPoint 350) were 18.4 grams, and one (Lotek PinPoint 450) was 19.1 grams. The weight of the kākā ranged from 444 grams to 588 grams and averaged 524 grams, resulting in the devices averaging 3.55% of the kākā’s body mass, with a range from 3.1% to 4.3%. These transmitter weights are all less than previous tracking studies of kākā that have observed no ill-effects (Powlesland et al., 2009; Recio et al., 2016). When possible, kākā were recaptured to remove the GPS device when the battery was depleted, which was expected to be 5-6 months given the GPS fix rate and duration of VHF and UHF windows, and health checks were again conducted. No tags detached via the weak link before the batteries depleted, and there were no observed adverse health effects due to the GPS devices or their attachment to kākā.

### Study site and semi-urban kaka reintroduction programs

Orokonui Ecosanctuary is a 307-ha sanctuary near the city of Dunedin (South Island, New Zealand) that is completely surrounded by a 1.9-metre-high pest-resistance fence. Since the time the Orokonui fence was erected in 2007, the reintroduction of multiple taxa has led to positive biodiversity impacts within and beyond the fence (Aichele et al., 2021; Bogisch et al., 2016; Jarvie et al., 2016; Tanentzap & Lloyd, 2017). The vegetation within Orokonui is predominately comprised of regenerating kanuka-broadleaf native forest with small areas of old growth forest, as well as stands of exotic pine forest (predominately *Pinus radiata*) and mountain ash (*Eucalyptus regnans*). Surrounding the sanctuary is a fragmented landscape of old-growth and regenerating native forest fragments, as well as exotic conifer forest, working farms for grazing livestock, and semi-urban areas. At the time of the study, adjacent areas outside the fence were subject to continuous pest-control, targeting mustelids, possums (*Trichosurus vulpecula*), and rodents. Supplementary food is available to kākā at feeding stations in Orokonui year-round as parrot-specific pellets and sugar water.

### Data collection

To gather locational data, we fitted SWIFT fix GPS devices (PinPoint GPS VHF 350/450, 18.4 g/19.1 g, Lotek, NZ) to 10 kākā, which were set to take a location every 3 hours for the nine PinPoint 350 devices, and every 2 hours for the single PinPoint 450 device due to a larger battery. The devices had VHF (very high frequency radio) for locating animals, and UHF (ultra-high frequency radio) for downloading data remotely. We had a 4-hour VHF window each day to locate the kākā, and an 8-hour UHF window each day to download the data. The UHF window was longer as it uses little battery and occasionally the data could be downloaded when individuals visited supplementary feeding stations, obviating the need for locating the individual via VHF. The GPS devices were distributed evenly between the sexes, with five females and five males, and across a range of ages from 1 – 10 years old (mean = 4.5 years). The tracked kākā were a combination of Orokonui-fledged individuals, which are more akin to wild birds and have little human intervention besides monitoring nest success, and captive-raised individuals, which are raised in an aviary breeding facility by adult breeding pairs until they are independent enough for soft-release into Orokonui. Kākā are considered dependent on adults for food until roughly 5 months of age (Moorhouse & Greene, 1995). All but one kākā had been in the sanctuary for a minimum of 5 months, with a range of time within the sanctuary of up to 10 years. The remaining kākā was a 10-year-old male who was released (with his breeding partner) into Orokonui directly from a captive-breeding facility, and the environment was therefore novel to this individual. The GPS devices recorded location data, the number of satellites, and horizontal dilution of precision (HDOP) on the 2- or 3-hour schedule listed above, as well as temperature and overall dynamic body acceleration (ODBA) on a separate schedule at 1-minute intervals for all devices. All GPS locational data was projected to New Zealand Transverse Mercator (NZTM2000 – EPSG:2193) before analysis. Prior to and during the study period (from late 2019 to early 2021) kākā mortalities of tracked and non-tracked individuals were recorded during surveys by managers of the sanctuary or by mortality signals from the GPS devices of this study or from VHF-only devices deployed separately to this study. Kākā were sent to Massey University, New Zealand, for post-mortem to determine cause of death, and we recorded the approximate time and location of the mortality where possible.

### Data Analysis

#### Location filtering

To remove unrealistic locations that are unlikely to represent the true location but are instead due by GPS-related error, we used a filtering method which is based on the maximum observed speed of each individual (Shimada et al., 2012, 2016). We calculated the maximum observed speed (Vmax) between successive locations for each individual, based on the locations from six or more satellites (average of 51.4% of locations were from ≥ 6 satellites per individual). As 95% of SWIFT-fix GPS locations are considered to be within 19.0m from 6 satellites (Forrest et al., 2022), we expected these observed maximum speeds to reflect the maximum speed dictated by the particular individual’s behaviour, rather than due to measurement error. The individual Vmax values were then applied to all locations for that individual, and locations that had a speed exceeding the Vmax before and after the location were removed from further analyses. The location filtering analysis was conducted using the R package ‘SDLfilter’ (Shimada et al., 2012).

#### Home range behaviour and presumed risk exposure

To estimate each individual’s home range, we used the weighted autocorrelated kernel density estimator (wAKDE) from the ‘ctmm’ package (Calabrese et al., 2016; Fleming et al., 2015; Silva et al., 2022). We used the perturbative Hybrid REML (pHREML) approach to estimate the best fitting movement process, although the results with maximum likelihood (ML) were very similar. The selected movement models were a combination of Ornstein-Uhlenbeck (OU) and Ornstein–Uhlenbeck Foraging (OUF) models, which are both appropriate for estimating range distributions (Alston et al., 2022; Calabrese et al., 2016). For the risk exposure analyses we used the entire probability density of the range distribution (denoted as home range - HR), as the HR quantifies the space that is expected to be used by individual over the longer term (Alston et al., 2022; Fleming et al., 2015). We can therefore sum cells of the discretised probability density to approximate the time the individual is expected to spend in a specified area. With this in mind, we calculated a proxy of presumed risk exposure by calculating the proportion of the HR that was outside the Orokonui Ecosanctuary fence, as the majority of threats to kākā (introduced predators and human-based threats such as infrastructure and vehicles) exist only outside of the fence.

#### Comparison of home range area between individuals

To compare home range sizes between individuals and groups, we considered the area contained within the 50% contour (HR_50_) to represent areas of core use, and within the 95% contour (HR_95_) as the home range (Börger et al., 2008; Kie et al., 2010; Kranstauber et al., 2012; Stark et al., 2017). To identify differences between individual’s space use, we assessed the influence of age, sex, and origin (Orokonui-fledged or captive-fledged) of individuals on home range area using a model-selection approach. We proposed candidate models including age, sex or origin as the covariates for generalised linear models (GLM) with a Gamma distribution and logarithmic link function to account for the non-negative, continuous and right-skewed data (Bartoń, 2020; Burnham et al., 2011). Due to the low sample size of 10 individuals, we did not consider multiple covariates in a single model. Models were ranked by the corrected Akaike information criterion (AICc) due to the small sample size (Johnson & Omland, 2004), and model fit was assessed using a likelihood-ratio based on pseudo-R-squared for each model (Bartoń, 2020). Dispersion (φ), Pearson residuals and quantiles were checked for model fit (Dunn & Smyth, 2014).

#### Dynamic space use

To assess how the space use of each individual changed throughout time we employed a ‘sweeping window’ framework. To achieve this we estimated occurrence distributions (ODs) using GPS locations from a temporal window of fixed width (e.g., a month of locations), which represents a snapshot of the animal’s space use. This snapshot is moved across the full study period incrementally (e.g., by a day), resulting in temporally overlapping ODs (Figure 1). We refer to the approach of temporally overlapping ODs as dynamic space use, as it allowed us to identify when and how space use changed with respect to measures such as OD area, positional shifts of the OD centroid, or OD overlap across temporal snapshots. Occurrence distributions are more appropriate than range distributions here, as ODs quantify the uncertainty within the animal’s movement path, therefore estimating the probability of finding an animal in a particular location during a specified time period, rather than estimating long-term home range (Alston et al., 2022; Fleming et al., 2015, 2016). The sweeping window for space use is similar to the approach of Schlägel et al. (2019), who estimated interactions between individual animals.

**Figure 1:**
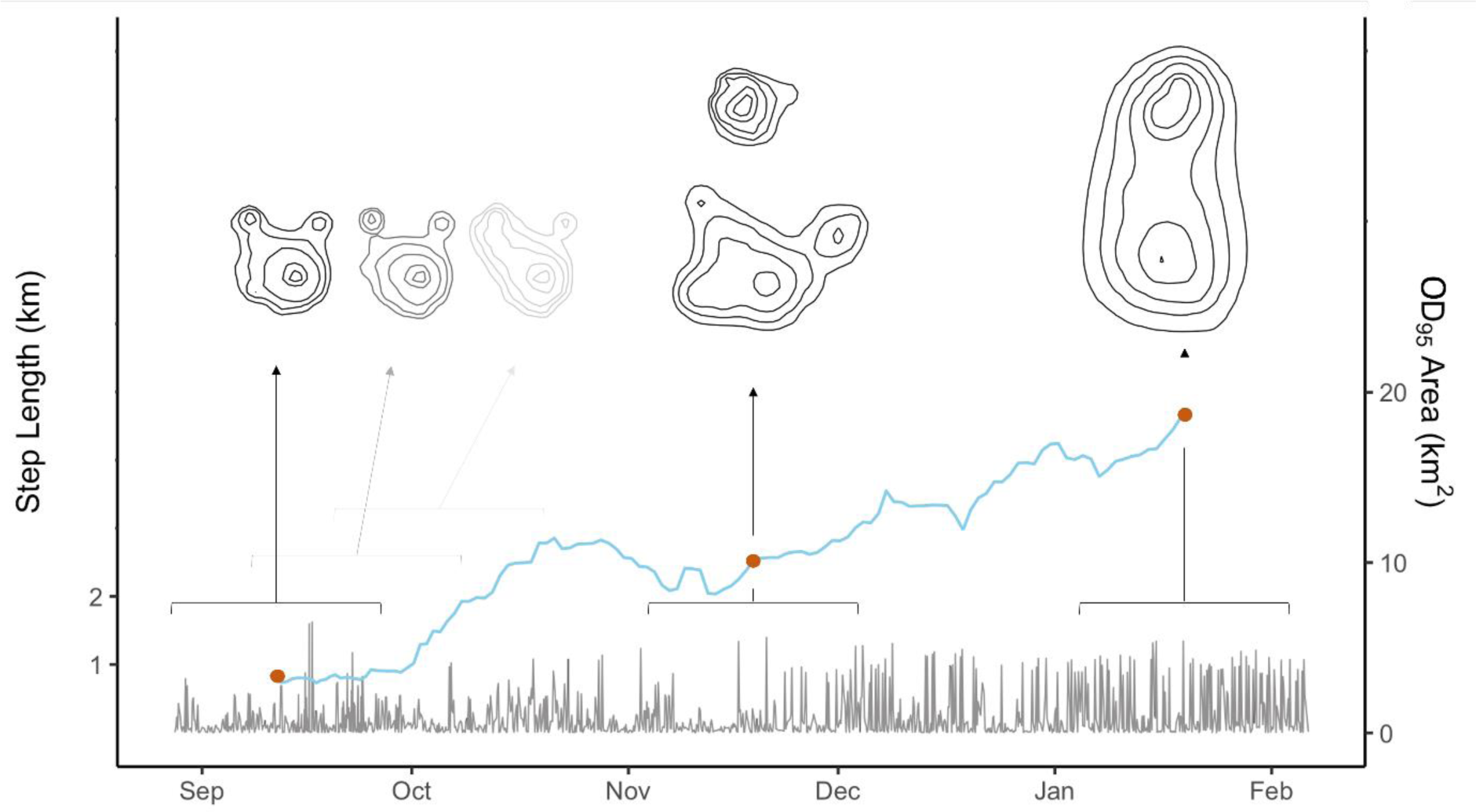
Concept plot of the dynamic space use method, based on data from this paper (kākā ID 08). Occurrence distributions (ODs) were estimated for temporally overlapping snapshots of a fixed window width of 240 locations, which was swept along the movement track at a fixed increment of 6 locations. For this individual it resulted in 169 temporally overlapping snapshots of space use. The kākā’s movement trajectory is represented by step lengths (black line – left y-axis), and the resulting OD_95_ areas that were calculated for each temporal snapshot are shown by the blue line (right y-axis). Interpretation of the area of occurrence distributions should be taken carefully however, as occurrence estimators only quantify uncertainty in the movement path, and can be sensitive to the sampling interval and idiosyncrasies of the animal’s movement behaviour (Alston et al., 2022). Several example ODs (all on the same scale) are shown to illustrate the gradual changes of space use. In this example, the large increases in OD_95_ were due to the inclusion of a new core area to the north in the central and right ODs. This is also reflected in the step lengths for the latter part of the tracking period, with a higher number of longer steps as this individual moved between the two core areas.

However, we focused on the dynamics of spatial behaviour more broadly, and calculated summary statistics such as the area, centroid, and overlap of each snapshot OD, which we tracked throughout time and compared between individuals. Interpretation of the area of occurrence distributions should be taken carefully however, as occurrence estimators only quantify uncertainty in the movement path and can be sensitive to the sampling interval and idiosyncrasies of the animal’s movement behaviour (Alston et al., 2022). A similar and useful approach outlining space use patterns for discrete, non-overlapping time periods is in Kranstauber et al. (2020).

Within the dynamic space use sweeping window framework there are two parameters, one that determines the number of locations in each temporal snapshot, which we call the window, and another we call the increment, which is the number of locations that the window is moved along the track by (Figure 1). We used a window containing 240 locations of the movement trajectory, which is approximately one month at 3-hour fix intervals, which was swept along at an increment of 6 locations, which is roughly 18 hours. The recorded timestamp of each OD was midway between the first and final locations. The window and increment widths can be freely chosen and should depend on the frequency of telemetry locations and questions being asked. Shorter window widths will capture changes in space use that are shorter in duration, but for identifying persistent changes in the pattern of space use the animal should ideally visit all of its home range during the window’s timespan. The only downside we see to a smaller increment is computation time, and it should be made as short as practicable. In this study, the fix-success rate (proportion of attempted GPS fixes that were successful) averaged 78.1% for all individuals, and therefore the actual window width will be wider than one month, and the increment may be greater than 18 hours. We kept the number of locations consistent rather than the time period to ensure that space use was estimated for a consistent number of locations. A consistent time period can also be used. In this case, we used dynamic Brownian bridge movement models (dBBMM) (Horne et al., 2007; Kranstauber et al., 2012) to estimate ODs separately for each temporal snapshot, although other occurrence estimators may also be used. The dBBMM method also requires the user to input a window and a margin size to calculate the likelihood, for which we used 15 and 3 locations, respectively. To prevent confusion, although the window and increment parameters of the dynamic space use framework are similar to the window and margin parameters of the dBBMM, they are separate, and both need to be specified by the user.

To assess the pattern of space use changes throughout the tracking period we constructed a similarity matrix of Bhattacharyya’s Affinity (BA), which represents the similarity between all the snapshot ODs for each individual (Fieberg & Kochanny, 2005; Kranstauber et al., 2020). This approach can be used to identify space use patterns, such as in Kranstauber et al. (2020). We opted for BA as it considers the probability density of the OD (which was discretised into cells), it is computationally efficient, and in this case all snapshot ODs within each individual at least partly overlapped. If each individual’s snapshot ODs did not overlap then earth mover’s distance may be more appropriate as it considers distance between non-overlapping ODs (Kranstauber et al., 2016). To quantify expansion and contraction of space use for each individual throughout time we calculated the OD_95_ area for each temporal snapshot, and to assess positional shifts in space use we calculated the location of the OD_95_ centroid. To compare positional changes of space use between individuals and groups, we calculated the distance between successive centroid locations and took the overall average for each individual, which we termed space use drift. The centroid for multimodal ODs was taken to be the overall OD_95_ centroid rather than the centroid of the largest OD_95_ area, which considers the inclusion of new areas of space use but also prevents artificial ‘jumps’ of the centroid if there were two modes of the OD that were similar in size but fluctuated as the largest in area. To statistically compare space use drift between kākā, we fitted a generalised linear model with a Gamma distribution and logarithmic link function and conducted the same model diagnostics as above.

## Results

### Data collection, data filtering and mortalities

The GPS unit data collection ranged from 111 to 168 days (mean ± SD = 147 ± 17 days), gathering between 725 to 1727 successful fixes for each individual (mean ± SD = 1087 ± 257 successful fixes). GPS fix-success rate of varied from 63.6% – 90.2% (mean ± SD = 78.1 ± 8.7%), resulting in a total of 10,868 successful location fixes for all individuals. The Vmax from 6 or more satellites ranged from 0.247 to 1.937 km/hr between individuals (mean ± SD = 0.948 ± 0.58 km/hr), resulting in between 0.23% to 3.29% (mean ± SD = 1.13 ± 0.93%) of locations being removed for each individual (Table S.1). During an 18-month period prior to and including the tracking period seven kākā were recorded to have died. Two deaths occurred prior to tracking and five deaths occurred during the study period - of these five, two were individuals that were not part of this study but were being radio tracked with VHF-only devices, and one was an individual that was part of this study. The GPS-tracked individual died within a month of deployment, so the data was excluded from the analyses in this paper due to the short tracking period, and the GPS device was then redeployed on another individual, and the data for that individual is included in this study. The four mortalities that were not being VHF- or GPS-tracked were discovered incidentally. Six of the kākā that died were of known age, and were all 3-years or younger with an average age of 1.8 years. Confirmed causes of mortality included brodifacoum poisoning and toxoplasmosis and suspected but unconfirmed causes included electrocution via powerlines (Table S.2). No mortalities or adverse effects were attributed to the GPS or other telemetry devices or the attachment harnesses.

### Static space use: home range area and comparison between individuals

Individual home range area (HR_95_ area) varied widely, from 0.34 km^2^ to 9.92 km^2^, with a mean ± SD = 4.12 ± 3.83 km^2^ (n = 10 individuals) (Figure 2 and Table S.1). The home ranges of all kākā were at least partly within the reserve, and three almost entirely, with the proportion of each kākā’s HR inside the Orokonui fence ranging from 0.11 to 0.98 (mean ± SD proportion of HR inside Orokonui = 0.59 ± 0.35) (Figure 2 and Figure 3B). There was no significant trend of HR_95_ area in relation to sex or origin (Table S.3 and Figure S.4) but there was a significant negative correlation between HR_95_ area and age (P = 0.001, R^2^ = 0.76 – Figure 3A). The average home range size for kākā 3 years and younger was 6.14 km^2^ (n = 6), whereas the average home range size for kākā 5 years and older was 1.09 km^2^ (n = 4) (Figure 2). Similar to home range area, the proportion of each individual’s HR that was outside of the Orokonui Ecosanctuary fence also decreased significantly with age (P < 0.001, R2 = 0.82 – Figure 3B).

**Figure 2:**
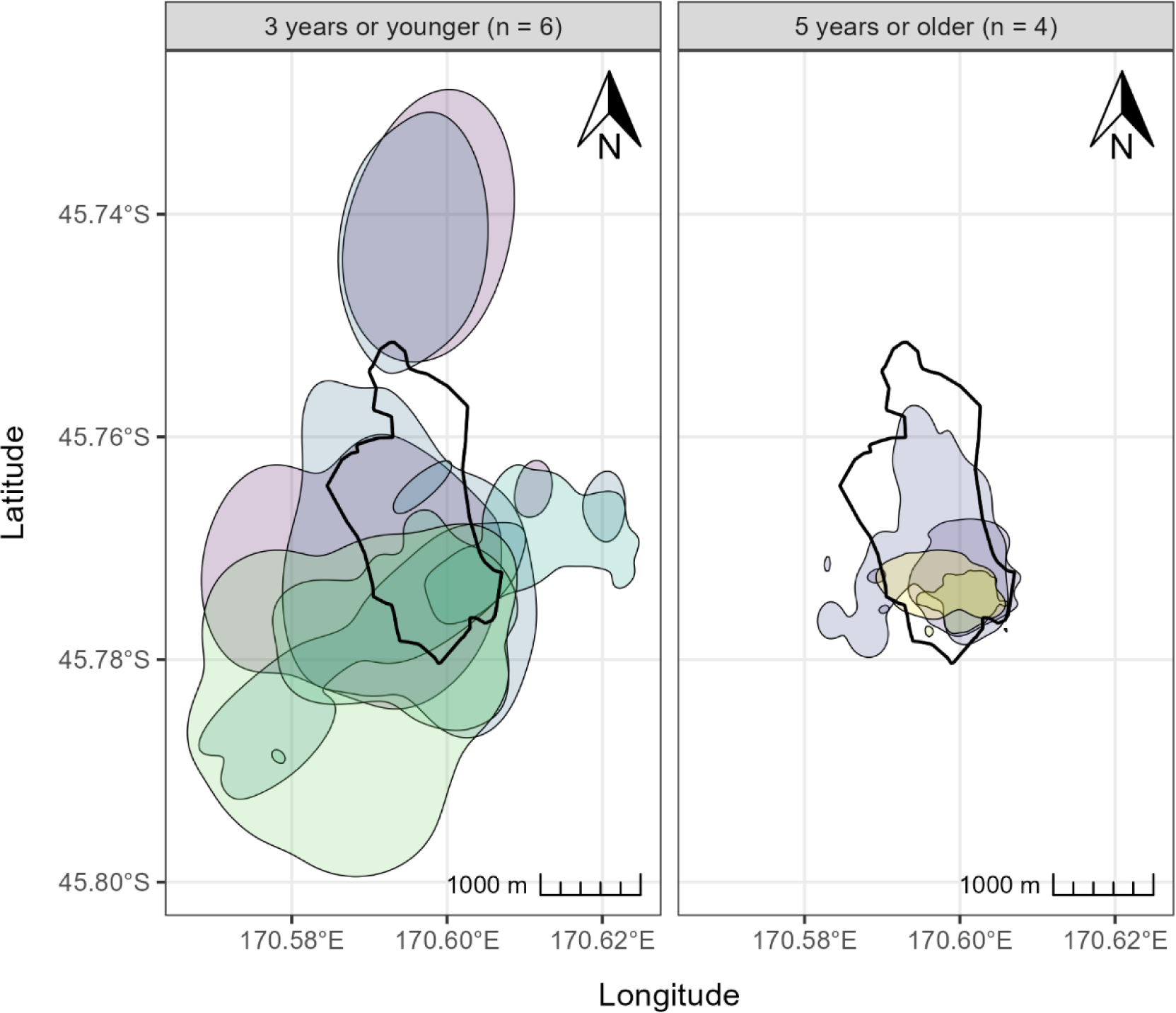
95% home range distributions (HR_95_) for each individual estimated from GPS tracking and weighted autocorrelated kernel density estimation (wAKDE). The left plot shows the HR_95_ isopleths for individuals 3 years or younger (n = 6), and the right plot shows the HR_95_ isopleths of individuals 5 years or older (n = 4). The fence of the Ōrokonui Ecosanctuary is shown as the thicker black line in the centre of each plot. The scale is the same for both plots to highlight the differences between HR_95_ areas of younger and older individuals.

**Figure 3:**
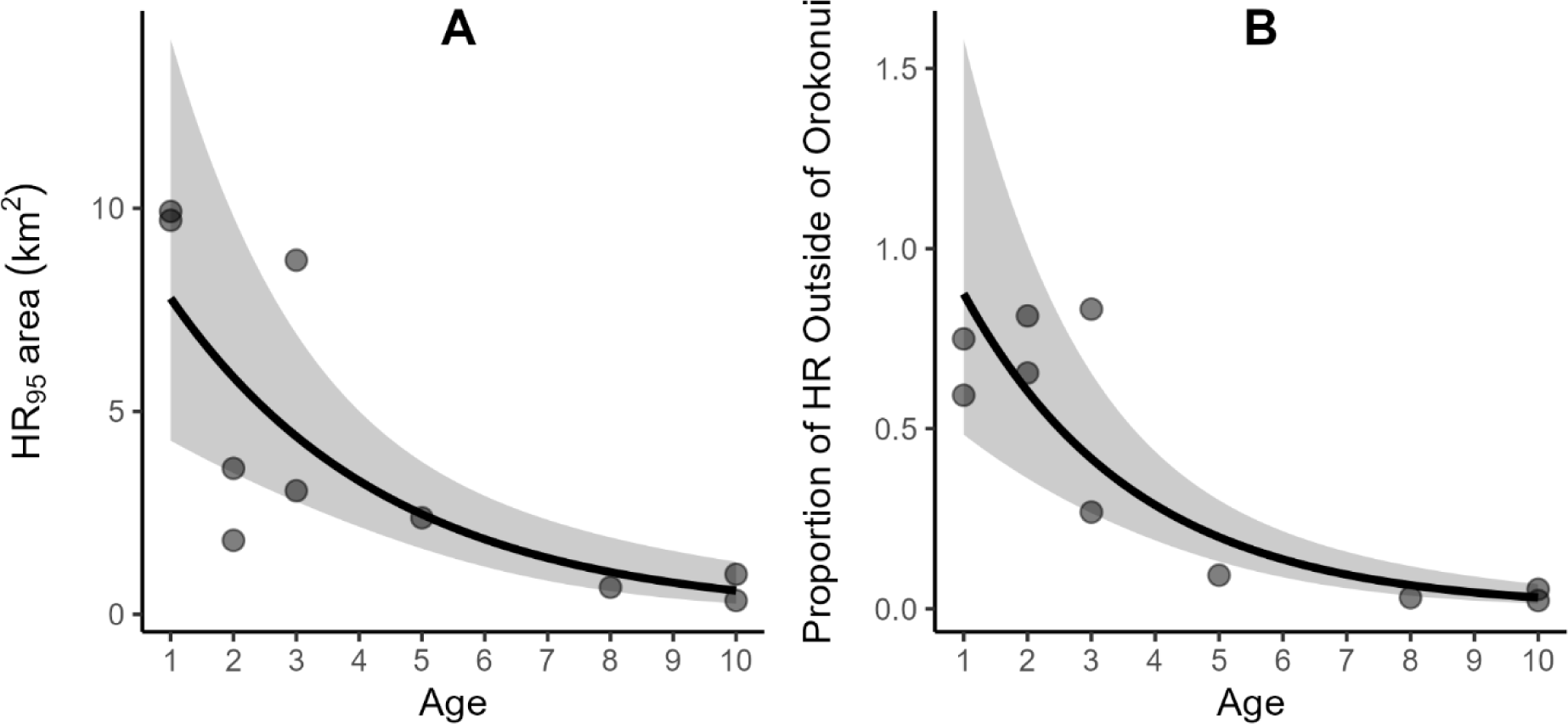
**A**) Home range area (HR_95_ area) as a function of age for 10 kākā GPS tracked at Ōrokonui Ecosanctuary, New Zealand. Generalised linear model using Gamma distribution with log link was fit with age as a predictor, ribbon is the 95% confidence interval. Pseudo-R^2^ based on likelihood ratio = 0.73. **B**) Proportion of individual home ranges (HR) of kākā that were outside of the Ōrokonui Ecosanctuary fence, as a function of kākā age. Generalised linear model using Gamma distribution with logarithmic link was fit with age as a predictor, ribbon is the 95% confidence interval. Pseudo-R^2^ based on likelihood ratio = 0.82.

### Dynamic space use

Space use was most variable for the two juvenile kākā (1-year old), with Bhattacharyya’s Affinity (BA) reaching as low as 0.43 and 0.38, whereas the minimum BA of all remaining individuals was 0.76 (Figure 4 and Figure S.5). The large dissimilarities in BA for the juvenile kākā were predominately due to positional changes of the OD_95_ centroid rather than changes in OD_95_ area. The area of every kākā’s OD_95_ varied throughout the study period, although the risk exposure through time remained largely constant for most individuals (Figure S.6 and S.7).

**Figure 4:**
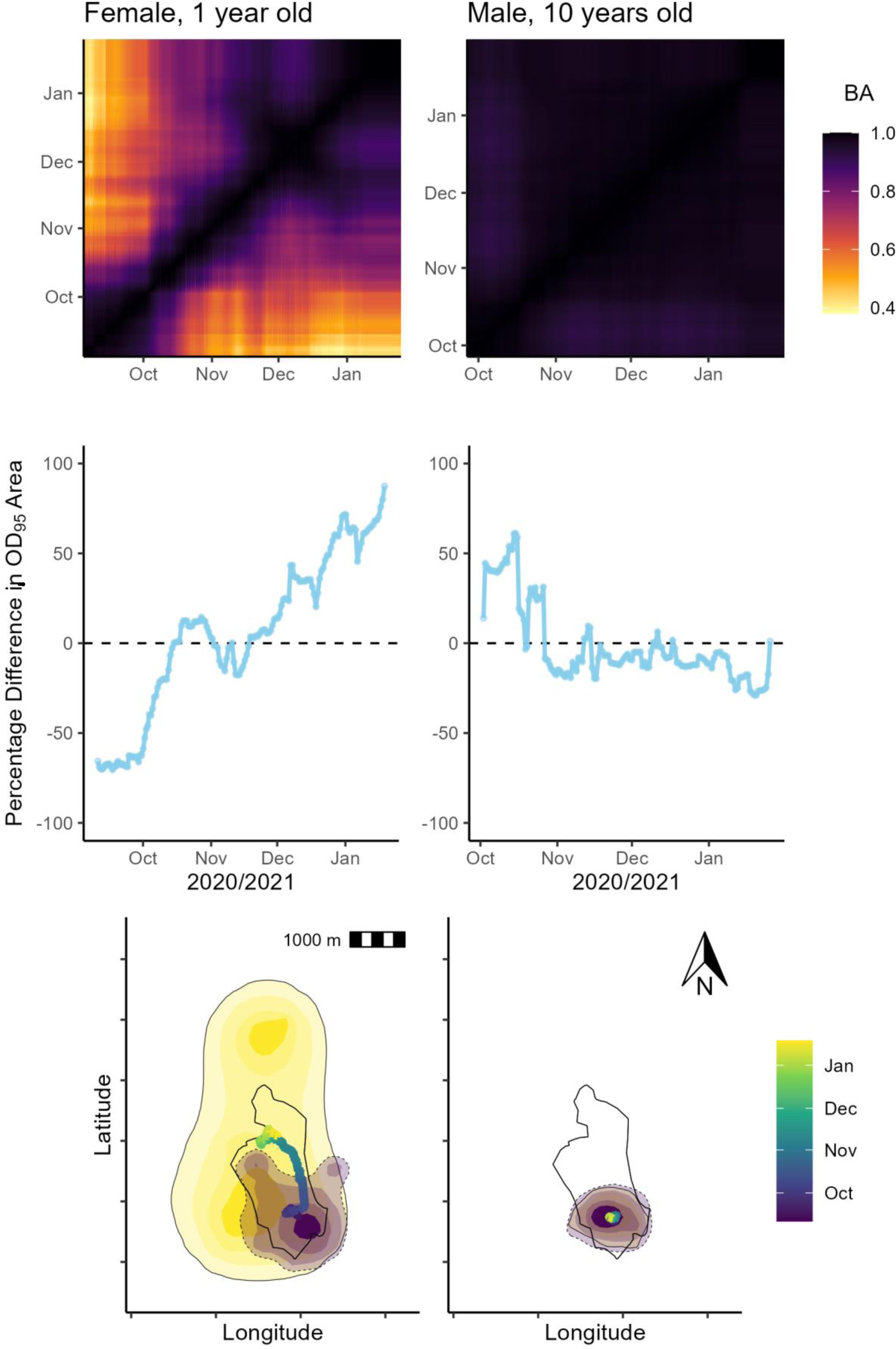
The left column represents the dynamic space use of an individual female kākā of age 1 year, and the right column represents the dynamic space use of a 10-year-old male kākā. The upper panels show the similarity matrices of Bhattacharyya’s Affinity (BA) throughout the study period. The middle panels show the percentage difference in OD_95_ area compared to each individual’s average OD_95_ area for the full study period. The lower panels show the drift of OD_95_ centroid on the same scale, with the colour relating to the time of year (purple to yellow), with the fence of Ōrokonui Ecosanctuary represented as the black outline. The lower panels also show the initial (purple with dashed line) and final (yellow with solid line) ODs for each individual. The scale bar is in 200m increments and is 1000m in total.

There was also some synchrony as to when OD_95_ area varied within the male and female groups, with females expanding their OD_95_ areas in September before contracting after October, and males expanding their OD_95_ areas after October (Figure S.6).

The centroids of the OD_95_ drifted farthest on average for the two juvenile kākā, with average drift distances ranging from 32.3 m to 4.87 m between temporally overlapping ODs (F_1,8_ = 38.52, P < 0.001, pseudo-R^2^ = 0.81 – Figure 5 and Figure S.8). There was no discernible population- level trend of space-use drift that might have related to seasonality (Figure 5B).

**Figure 5:**
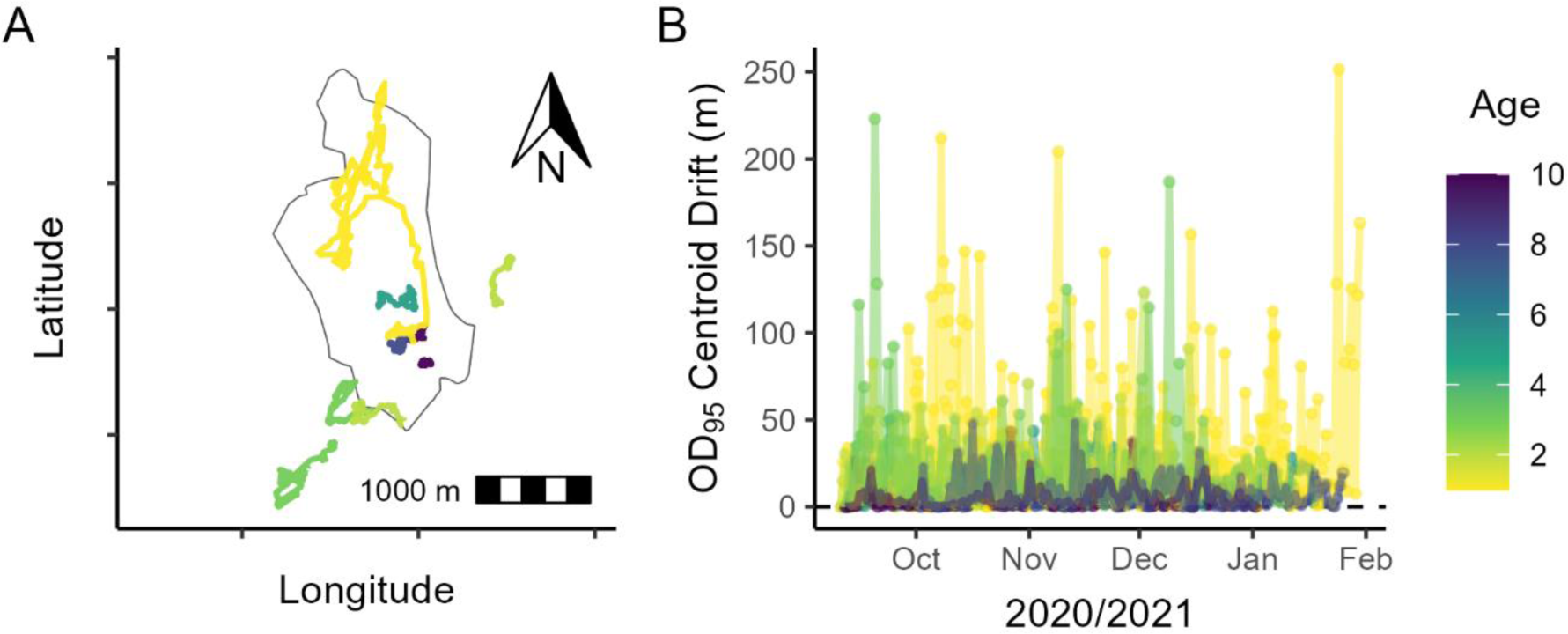
**A**) Position of the UD_95_ centroid for each individual kākā throughout the study period, in relation to age (colour – legend to the right). The fence of Ōrokonui Ecosanctuary is shown in black. The scale bar is in 200m increments and is 1000m in total. **B**) Distance between UD_95_ centroids from one space use snapshot to the next. There appear to be no discernible temporal trends in space use drift.

## Discussion

In this study, we used home range (HR) and dynamic space use analyses using temporally overlapping occurrence distributions (ODs) to assess risk exposure in a reintroduced kākā (*Nestor meridionalis*) population. Summarily, the HRs of younger individuals (less than 3 years) were much larger than that of older individuals, and the space use of juvenile kākā (1-year-old) was more temporally variable, with significant shifts of the OD_95_ centroid. We therefore suspect that the larger home ranges and more exploratory behaviour of younger individuals put these cohorts at higher risk than older individuals due to more time spent outside the reserve. These findings correlate with higher incidental mortality observations of younger individuals, of which the average known age was 1.8 years. All kākā included the supplementary feeding stations within the core area of their HR, which suggests there may be a reliance on supplementary feeding stations that provides an anchoring effect, which may help to increase the safety of kākā by reducing their time spent in risky areas. Additionally, as we caught individuals within the sanctuary, it is possible there are kākā that have home ranges exclusively outside the sanctuary that we did not catch to attach GPS devices to.

The age-related differences in home range area and dynamics may be due to several factors. Namely, 1) younger kākā may be more exploratory; 2) there may be intraspecific conflict between age groups leading to displacement; 3) there may be a selection pressure against large home ranges; and 4) familiarity rather than age may explain the trend. For 1), as shown in previous studies of wild kākā and other bird species, older individuals are typically more neophobic and more efficient foragers, which can reduce the need for large home ranges relative to younger individuals (Bond & Diamond, 2004; Clay et al., 2018; Loepelt et al., 2016; Sherratt & Morand-Ferron, 2018). This trend may also be compounded by the constant availability of supplementary food from feeding stations, as they needn’t forage widely. The more dynamic space use in juvenile kākā predominately arose from positional changes of the OD_95_ centroid, which was due to the initial visitation of novel areas that were thereafter visited regularly, which suggests exploratory behaviour followed by exploitation (Berger-Tal et al., 2014). This pattern may also indicate information gathering that may precede dispersal from their natal area (Andreassen et al., 2002; Ronce, 2007), and would provide valuable evidence towards home range establishment, which is difficult to test empirically (although see Maor- Cohen et al. (2021)). 2) it is also possible there is agonistic behaviour (intraspecific conflict), where older individuals are more dominant in areas with supplementary feeding stations, which displaces the younger individuals to the periphery, necessitating exploration of novel areas. However, we did not visually observe agonistic behaviour, all individuals still included the feeding stations in their core home ranges, and previous work has suggested that juvenile kākā at Orokonui visit the feeding stations regularly (Aichele et al., 2021). This evidence suggests that there isn’t strong displacement by older kākā against younger kākā, at least at feeding stations. 3) another possibility is that individuals that have survived to adulthood have always had small home ranges, suggesting a negative selection pressure against large home ranges. Definitively addressing this question would require tracking for longer to assess whether home ranges begin to contract in younger individuals. From our data, younger individuals had consistently larger home ranges, which declined uniformly, rather than younger individuals having both small and large home ranges, which would undergo a filtering process against large home ranges as age increased (Class et al., 2019). 4) we cannot definitively separate age and familiarity, which are inextricably linked but have relevance for conservation translocations, as ages may differ despite the environment being novel to all. Therefore, it is possible in our case that older individuals have smaller home ranges because they are more familiar with the area, and this is likely to be at least partly the case. However, the smallest home range in our sample was of a 10-year-old kākā that was released into the sanctuary at the time of tracking, so the area was completely novel to this individual. It should be noted though that this individual was raised in captivity, which has likely led to a reliance on supplementary food, and results may differ for wild adults translocated into the sanctuary. Summarily, as younger kākā still access supplementary food, the trend of larger and more dynamic space use in younger individuals appears to be predominately explained by exploratory behaviour, which is likely linked to familiarity. The exploratory behaviour is possibly preceding long-term home range establishment, which would be indicated by a contraction and stability of space use. If that is the case, it is unknown whether the new, smaller home ranges would be inside or outside of the sanctuary, as there were areas of core use (HR_50_) for several younger kākā outside the sanctuary.

Despite clear trends in the age-related difference in home range area and patterns of dynamic space use, we acknowledge the sample size of ten individuals from a single study site may not represent the full breadth of behavioural variability present in this or in other populations, and tracking more individuals may reveal more conclusive or even contradictory evidence. Given our timeline and funding, we prioritised fewer devices with a relatively high frequency of locations (i.e., GPS) to understand fine-scale space use, rather than solely VHF radiotracking devices that are less expensive and last longer, but do not have the consistency or frequency of locations to answer the questions we have addressed here. The duration of tracking also did not cover a full year and we may have missed seasonal variation outside of the tracking period.

However, the coverage of sex and age groups provided valuable and convincing evidence towards age-related differences in spatial behaviour, and the relatively fine-scale tracking data was important to provide sufficient resolution to estimate dynamic space use.

Although at least one other method has been developed to segment an animal’s trajectory with a sweeping window framework (Schlägel et al., 2019), how an animal’s space use changes gradually throughout time and whether this aligns with certain groups has otherwise been given little attention in the literature. We hope to address this by providing an easily applicable and intuitive dynamic space use estimation framework, and we highlight that different statistical space use estimators can be substituted for the dBBMM occurrence estimator, such as the generalized time-series Kriging approach which can accommodate multiple movement models (Fleming et al., 2016). Estimating both home ranges using a range estimator and dynamic space use using an occurrence estimator allowed us to strengthen our understanding of the kākā’s spatial behaviours by assessing total space used and how that changed throughout time. Tracking the area, centroid and overlap of the occurrence distributions allows for space use patterns to be identified as the ODs expand, contract, and shift in position, which can allow a multitude of questions about site fidelity, space use patterns, dispersal and exploration to be posed (Börger et al., 2008; Kranstauber et al., 2020, 2016).

Differences in spatial behaviour could lead to increased risk for certain groups of taxa in conservation reserves, particularly when threats remain in the wider landscape. In this study, the exposure to presumed risk did not vary substantially within groups throughout time, but this may not be the case if the kākā were tracked for longer, or for other taxa that may have more predictable seasonal behavioural changes. Species that increase their space use area or decrease their site fidelity to develop sufficient condition for breeding or hibernation, or to take advantage of ephemeral resources, may be exposed to greater risk as they venture beyond safe regions. Understanding which groups or individuals of a species are at the highest risk may assist managements decisions, particularly when planning translocations or reintroductions.

## Management Implications

Management of birds and other species of wide-ranging behaviour in conservation reserves should consider differences between groups of individuals that may lead to differential exposure to risk beyond areas of safety. This will also be relevant to the planning of translocations and reintroductions, as individuals are likely to differ in their spatial behaviours (Stuber et al., 2022), which may relate to age. In that case it may be fortuitous to translocate individuals from a range of ages if possible, although there are many factors that contribute to successful translocations, which may vary by species (Miskelly & Powlesland, 2013; Spatz et al., 2023). At the very least it should be considered that animals of varying ages may respond differently to novel environments, and younger individuals may be more likely to explore spatially. In fragmented landscapes that contain atypical features such as urban areas, infrastructure and exotic predatory species, reintroduced animal species will need to explore and adapt to survive, which may lead to innovations and a greater behavioural repertoire, improving their acquisition of resources and avoidance of threats. This study additionally highlights the key importance of appropriate management of threats outside of conservation reserves, particularly for wide-ranging flighted species. For other risk-exposed taxa, we recommend quantifying both home ranges and dynamic space use to assess and understand the area requirements and space use patterns of individuals, and to determine when and where animals may be exposed to risk. We provide code and data such that all the analyses and figures in this study can be reproduced.

