## Supplementary Figures and Tables for "Home range and dynamic space use reveals age-related differences in risk exposure for reintroduced parrots"

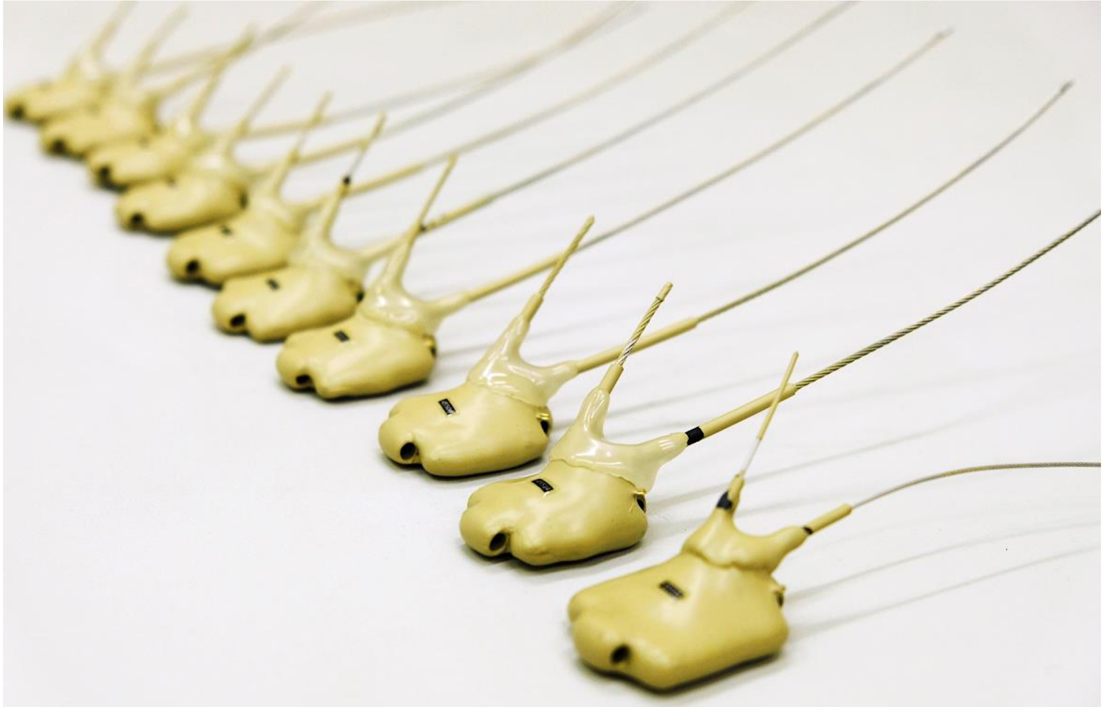

*Figure S. 1: The ten Lotek GPS devices (18.4 – 19.1 grams) that were used to collect GPS location data of kākā. These devices used a SWIFT fix algorithm. Smaller antenna is for connection to satellites, and longer antenna is for VHF and UHF communication.*

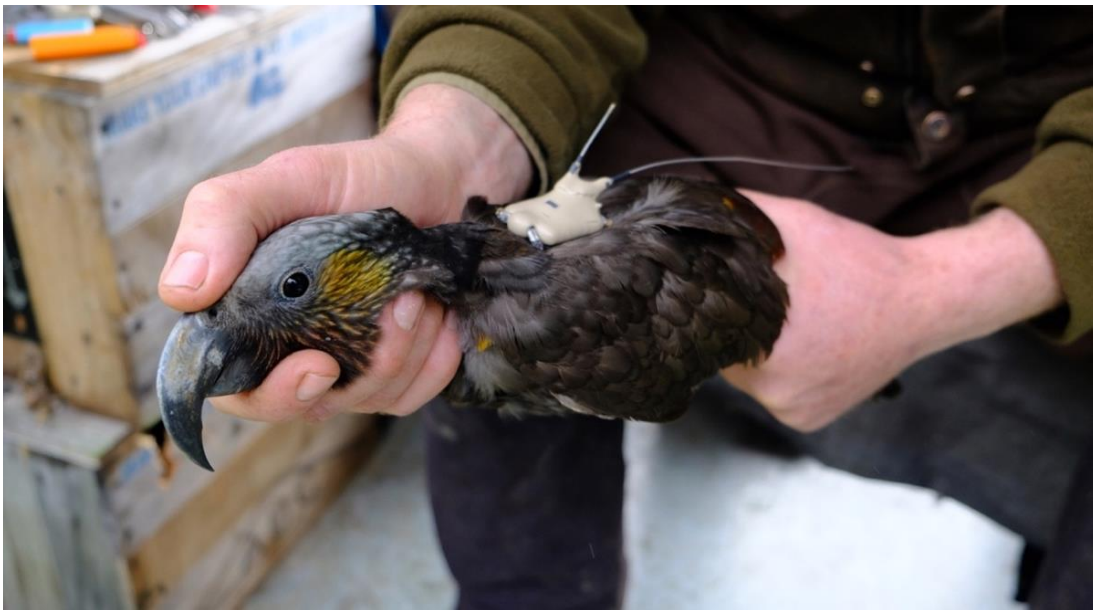

*Figure S. 2: Kākā with GPS device fitted using a backpack harness with a weak-link (Karl & Clout, 1987).*

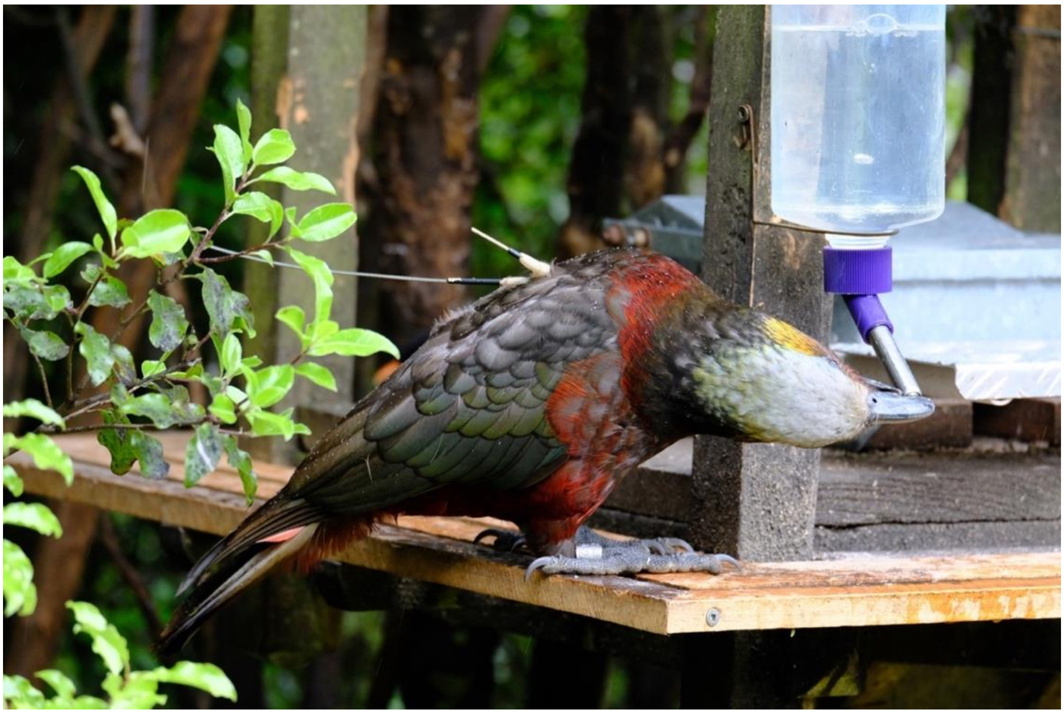

*Figure S. 3: Kākā with GPS device in-situ at a supplementary feeding station. Smaller antenna is for connection to satellites for GPS, and longer antenna is for VHF and UHF communication.*

*Table S. 1: Performance of GPS units and home range estimates of 10 kākā at Orokonui Ecosanctuary. Origin is either Orokonui-fledged (wild nest) or captive-raised in a breeding facility. Days – number of GPS data collection days; n – number of successful fixes; FSR – fix-success rate (proportion of successful fixes / failed fixes; filtered – percentage of data that was removed by maximum speed filter. $HR_{50}$ is the 50% isopleth of the range distribution estimated from autocorrelated kernel density estimation (AKDE), which can be denoted as the high use or core area; $HR_{95}$ is the 95% isopleth, which can be denoted as the full home range; $HR_{95}$ outside Orokonui is the proportion of $HR_{95}$ outside the fence of Orokonui Ecosanctuary.*

| Kākā | | | | GPS Unit Performance | | | | | | | Isopleth Areas | |
|---|---|---|---|---|---|---|---|---|---|---|---|---|
| ID | Sex | Age | Origin | Days | n | FSR | Vmax (km/h) | Filtered (%) | $HR_{50}$ ($km^2$) | $HR_{95}$ ($km^2$) | HR Inside Ōrokonui | HR Outside Ōrokonui |
| 05 | M | 1 | Orokonui | 140 | 1727 | 0.84 | 1.92 | 0.23 | 1.86 | 9.92 | 0.25 | 0.75 |
| 06 | M | 10 | Orokonui | 148 | 1003 | 0.78 | 0.45 | 1.69 | 0.14 | 0.99 | 0.95 | 0.05 |
| 07 | M | 5 | Captive | 162 | 1128 | 0.81 | 0.66 | 1.42 | 0.25 | 2.38 | 0.11 | 0.09 |
| 08 | F | 1 | Orokonui | 163 | 1257 | 0.90 | 1.20 | 0.32 | 1.90 | 9.71 | 0.41 | 0.59 |
| 09 | F | 3 | Orokonui | 128 | 725 | 0.64 | 0.83 | 0.55 | 0.24 | 3.04 | 0.63 | 0.27 |
| 10 | F | 2 | Orokonui | 111 | 828 | 0.73 | 0.97 | 1.21 | 0.19 | 1.82 | 0.19 | 0.81 |
| 11 | F | 2 | Orokonui | 168 | 916 | 0.64 | 0.88 | 0.44 | 0.70 | 3.60 | 0.35 | 0.65 |
| 12 | M | 3 | Captive | 157 | 1108 | 0.83 | 1.91 | 0.63 | 2.21 | 8.72 | 0.17 | 0.83 |
| 13 | M | 10 | Captive | 149 | 1034 | 0.80 | 0.23 | 3.29 | 0.03 | 0.34 | 0.98 | 0.02 |
| 14 | F | 8 | Orokonui | 143 | 1142 | 0.83 | 0.42 | 1.49 | 0.08 | 0.67 | 0.97 | 0.03 |
| Mean | | **4.5** | | **147** | **1087** | **0.78** | **0.95** | **1.13** | **0.76** | **4.12** | **0.59** | **0.41** |
| ± SD | | **3.6** | | **17** | **275** | **0.09** | **0.58** | **0.93** | **0.87** | **3.83** | **0.35** | **0.35** |

*Table S. 2: Kākā mortalities of the Orokonui kākā observed in the previous 18 months. Transmitters were different devices; Very High Frequency (VHF) transmitters were used for survivorship and nest monitoring and Global Positioning System (GPS) devices were used for the present study.*

Kākā Mortalities

| Approximate Date | Transmitter? | Age | Sex | Origin | Cause |
|---|---|---|---|---|---|
| November 2019 | - | 3 | F | Orokonui | Brodifacoum poisoning |
| November 2019 | - | 2 | F | Orokonui | Brodifacoum poisoning |
| July 2020 | - | - | - | - | Unconfirmed |
| August 2020 | VHF | < 1 | F | Captive | Electrocution (Powerline) |
| August 2020 | - | 3 | M | Captive | Toxoplasmosis |
| September 2020 | GPS | < 1 | M | Captive | Electrocution (Powerline) |
| December 2020 | VHF | < 1 | M | Captive | Unconfirmed |

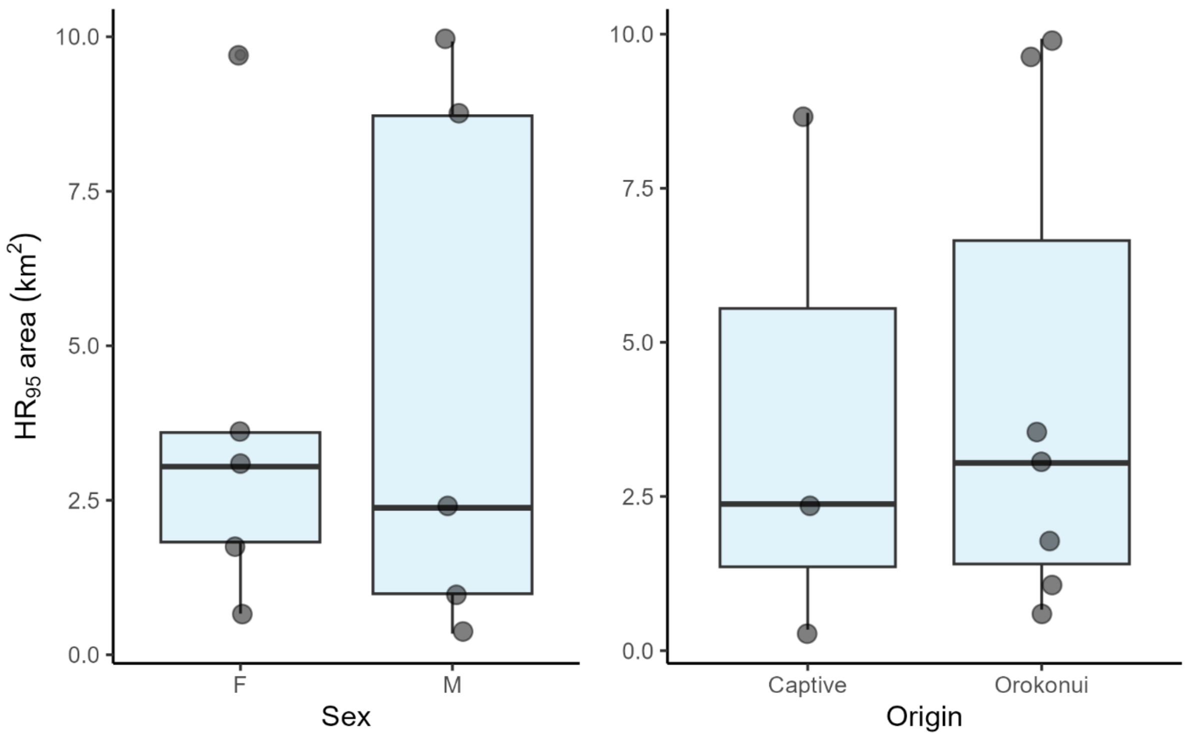


*Figure S. 4: Boxplots of the $UD_{95}$ areas for categories of sexes and origins of kākā that were GPS tracked out of Orokonui Ecosanctuary. Boxes indicate 25%, 50% (median) and 75% quartiles.*

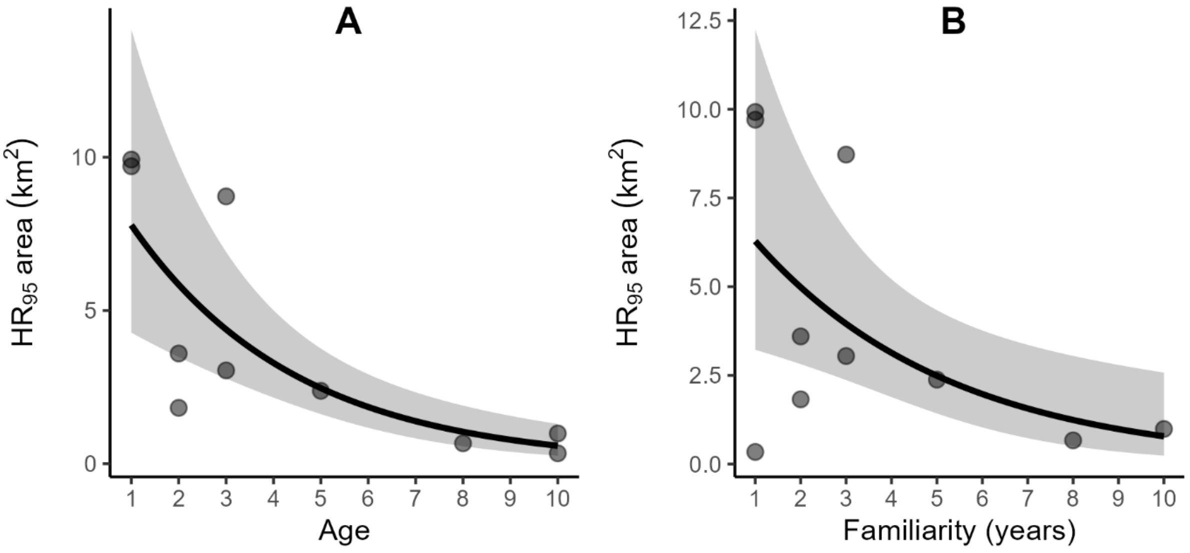


*Figure S. 5: Figure 1:* ***A****) Home range area ($HR_{95}$ area) as a function of age for 10 kākā GPS tracked at Ōrokonui Ecosanctuary, New Zealand. Generalised linear model using Gamma distribution with log link was fit with age as a predictor, ribbon is the 95% confidence interval. Pseudo-$R^2$ based on likelihood ratio = 0.73.* ***B****) Home range area ($HR_{95}$ area) as a function of familiarity for the same 10 kākā. Generalised linear model using Gamma distribution with log link was fit with familiarity as a predictor, ribbon is the 95% confidence interval. Pseudo-$R^2$ based on likelihood ratio = 0.36.*

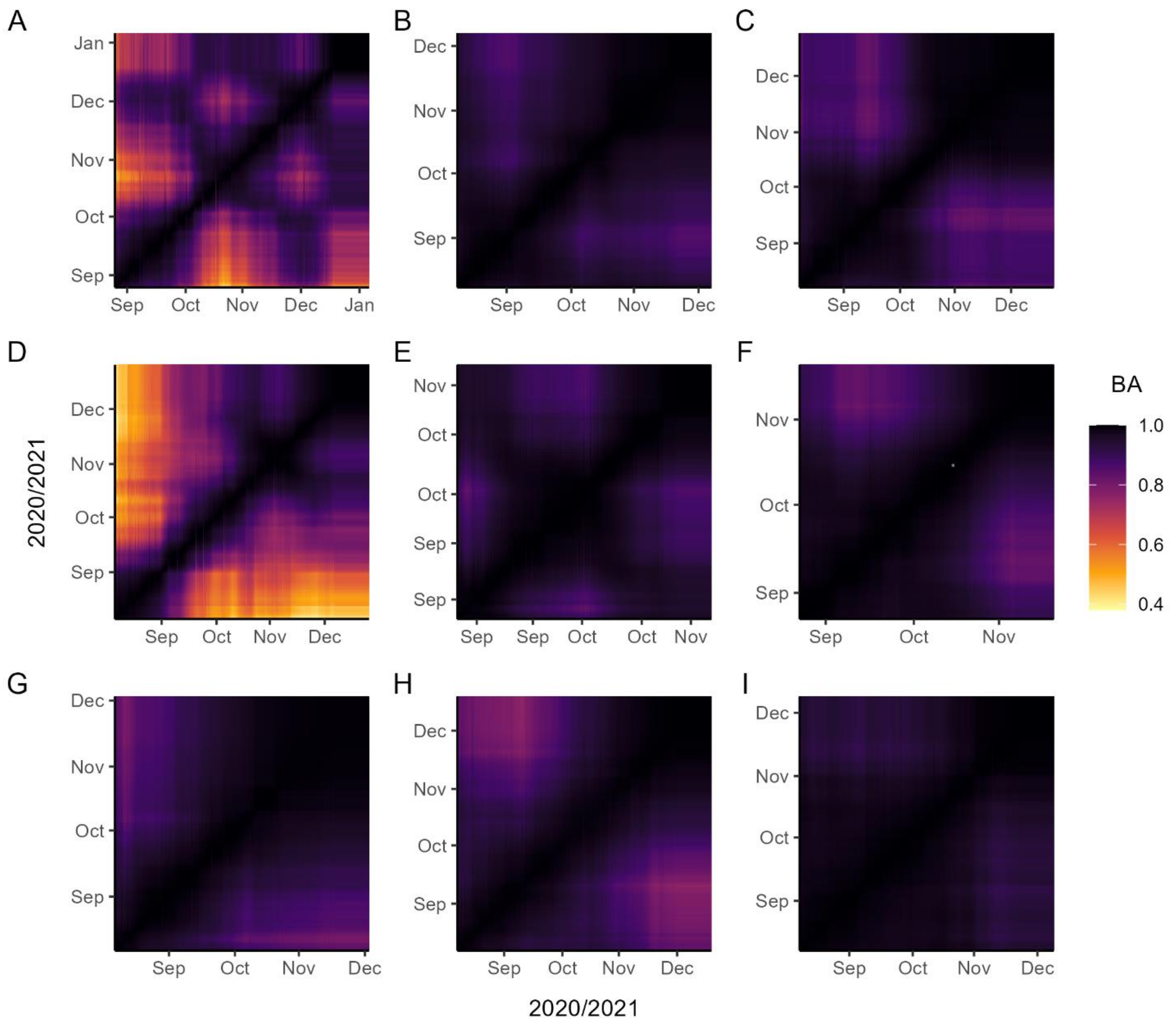


| Plot | A | B | C | D | E | F | G | H | I |
|---|---|---|---|---|---|---|---|---|---|
| ID | 45505 | 45506 | 45507 | 45508 | 45509 | 45510 | 45511 | 45512 | 45513 |
| Age | 1 | 10 | 5 | 1 | 3 | 2 | 2 | 3 | 10 |
| Sex | M | M | M | F | F | F | F | M | M |

*Figure S. 6: Similarity matrices of space use calculated with Bhattacharyya's Affinity (BA) between all UDs for each individual throughout the study period. Darker values indicate higher similarity and lighter values indicate lower similarity. The brighter colours of panels A and D indicate the dissimilarity between the UDs of the juvenile individuals, which was largely due to positional changes of the UD, rather than expansion and contraction.*

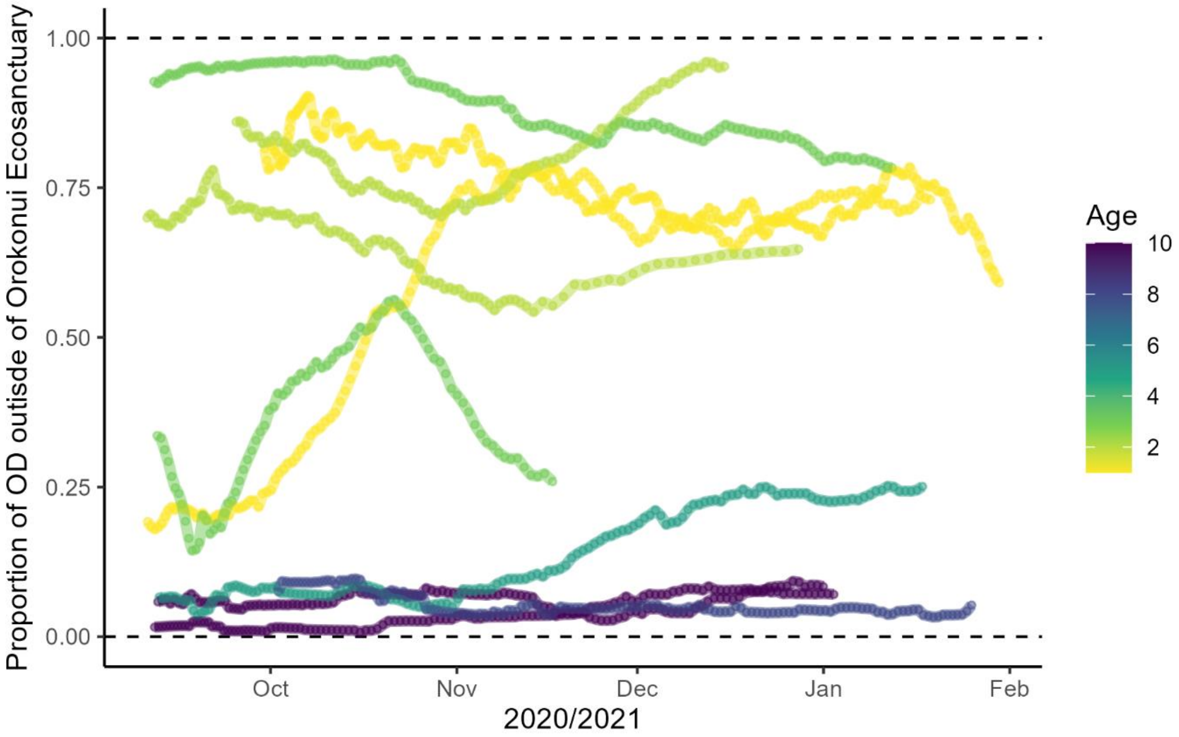


*Figure S. 7: Proportion of each snapshot utilisation distribution of each individual that was outside of the Orokonui Ecosanctuary fence, coloured by the age of the kākā. The space use of older individual's was predominately inside the fence, in an area of low presumed risk, which was relatively stable throughout time. Younger individuals had less stable risk-exposure throughout the study period, although there is no discernible trend between individuals that may indicate a period of heightened risk for the population.*

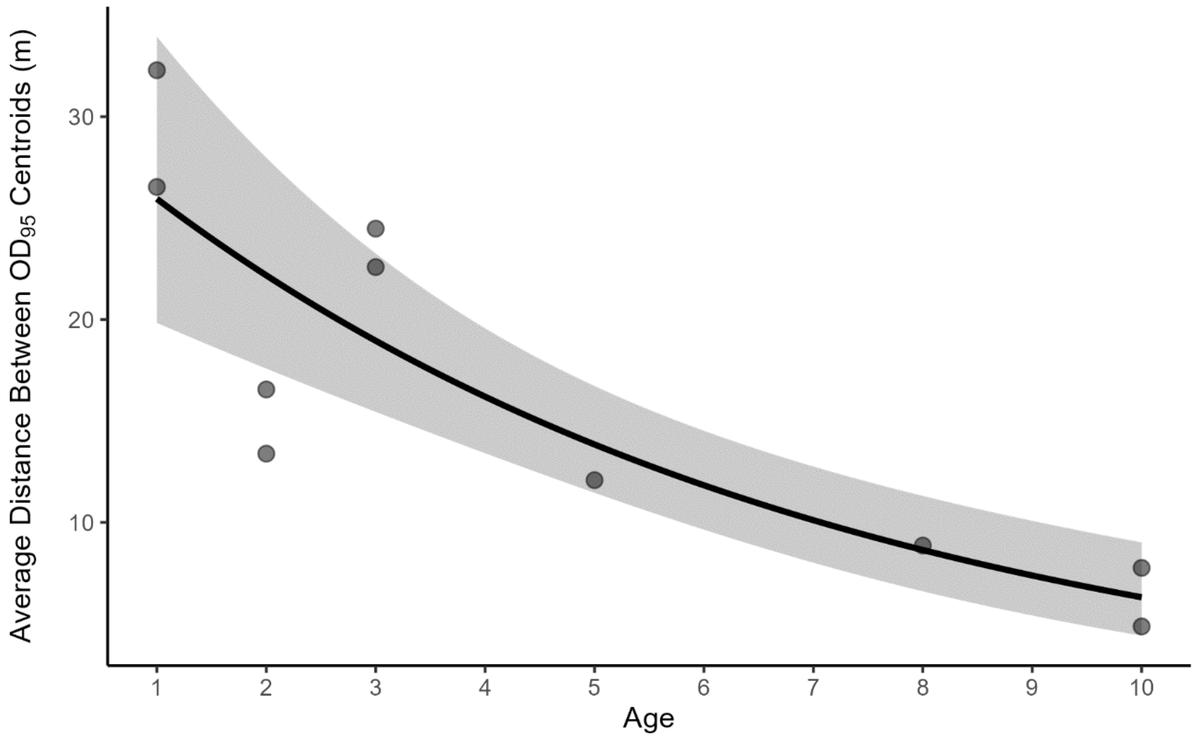


*Figure S. 8: Average distance between centroids of consecutive $UD_{95}$ snapshots from the sweeping window approach to quantify space use variability, which is plotted as a function of age. The increments between space use estimations were approximately one day. Larger values indicate that the $UD_{95}$ centroid changed position more than for smaller values, suggesting positional changes of space use. Line estimated from generalised linear model using Gamma distribution with logarithmic link, which was fitted with age as a predictor. The ribbon is the 95% confidence interval. Pseudo-$R^2$ based on likelihood ratio = 0.81.*

Alston, J. M., Fleming, C. H., Noonan, M. J., Tucker, M. A., Silva, I., Folta, C., Akre, T. S. B., Ali, A. H., Belant, J. L., Beyer, D., Blaum, N., Böhning-Gaese, K., de Paula, R. C., Dekker, J., Drescher-Lehman, J., Farwig, N., Fichtel, C., Fischer, C., Ford, A. T., … Calabrese, J. M. (2022). Clarifying space use concepts in ecology: range vs. occurrence distributions. In *bioRxiv* (p. 2022.09.29.509951). https://doi.org/10.1101/2022.09.29.509951
